# APOBEC3 activity and polymerase-ε deficiency are associated with distinct *IDH1* R132 hotspot mutations

**DOI:** 10.64898/2026.07.10.737816

**Authors:** Kelly E. Butler, Bilal Lone, Ecem Unal, A. Rouf Banday

**Affiliations:** Genitourinary Malignancies Branch, Center for Cancer Research, National Cancer Institute, National Institutes of Health, Bethesda, MD, 20892

**Author notes:** Correspondence: A. Rouf Banday, PhD Center for Cancer Research National Cancer Institute, NIH 37 Convent Dr Rm 1066A Bethesda MD 20892 E. Abbreviations: acute myeloid leukemia (LAML), Apolipoprotein B mRNA-editing enzyme, catalytic polypeptide threes (APOBEC3s), cholangiocarcinoma (CHOL), glioblastoma (GBM), Isocitrate dehydrogenase 1 (IDH1), lower grade glioma (LGG), replication fork directionality (RFD), skin cutaneous melanoma (SKCM), uracil DNA glycosylase (UDG).

**Keywords:** cancer, genetics, IDH1, mutational signatures, APOBEC3s, DNA polymerase

## Abstract

*IDH1* R132 mutations are among the most frequent hotspot mutations in cancer, but their mutational origins have remained unclear. Here, we provide evidence that *IDH1* R132C, the predominant *IDH1* mutation in cholangiocarcinoma, acute myeloid leukemia, and melanoma, likely arises through APOBEC3-mediated mutagenesis. *IDH1* R132C is a TpC>TpT substitution on the lagging-strand DNA template within a hairpin-forming sequence context, consistent with APOBEC3 susceptibility. *In vitro* assays showed that APOBEC3A can deaminate the relevant cytosine, and *APOBEC3A* and *APOBEC3B* were relatively highly expressed in tumor types with recurrent *IDH1* R132C mutations. *IDH1* R132G, a TpC>TpG substitution at the same site, may similarly result from APOBEC3 activity. By contrast, *IDH1* R132H, the predominant *IDH1* mutation in lower grade glioma and glioblastoma, is a CpG>TpG substitution at a methylated cytosine on the leading-strand DNA template, a pattern more consistent with DNA polymerase epsilon replication error. Concordantly, tumor types enriched for *IDH1* R132H showed relatively low *POLE* expression. Together, these in vitro and bioinformatic analyses provide insight into the distinct mutational mechanisms that likely underlie recurrent *IDH1* hotspot mutations in cancer.

## Introduction

Numerous distinct mutational processes contribute to carcinogenesis and leave characteristic mutational signatures, defined by the base substitution and surrounding sequence context (1). One of the most prevalent endogenous mutational processes in cancer is mediated by apolipoprotein B mRNA-editing enzyme catalytic polypeptide-like threes (APOBEC3s), which are cytosine deaminases that facilitate innate antiviral defense (2). However, APOBEC3s are frequently overexpressed in cancer and can deaminate host DNA, thereby generating oncogenic mutations (2). APOBEC3s preferentially deaminate single-stranded DNA, such as within hairpin loops and lagging strand DNA templates during replication (3–6). APOBEC3-mediated cytosine deamination generates uracil, and subsequent erroneous repair or replication of the uracil-containing template generates C>T or C>G mutations (2). These mutations underlie the COSMIC single-base substitution signatures SBS2 and SBS13, respectively (7). Among the seven APOBEC3s, APOBEC3A and APOBEC3B are most strongly linked with SBS2 and SBS13 (2, 7). While TCW (W = A or T) is common, APOBEC3s can also deaminate at TCG and TCC motifs including driver mutations (8–11).

Another common mutational process that generates C>T substitutions is spontaneous deamination of 5-methylcytosine (7, 12). This process is tightly linked with aging and classically represented by the “clock-like” signature SBS1 (7, 12). However, recent studies have identified DNA polymerase epsilon error as an additional potential source of C>T mutations at methylated CpG sites (13, 14). DNA polymerase epsilon carries out leading-strand DNA synthesis, and erroneous incorporation of adenine opposite 5-methylcytosine can generate C>T transitions independent of other polymerases or repair pathways (13, 14). The proofreading fidelity of DNA polymerase epsilon decreases with template-strand methylation, making replication errors particularly frequent at CpG sites (13, 14). DNA polymerase epsilon exonuclease domain mutations are canonically associated with SBS10 and SBS14 (1, 7). Wild-type DNA polymerase epsilon may also generate CpG>TpG mutations and contribute to additional mutational signatures, although its error rate is substantially lower than that of mutant polymerase epsilon (13, 14). Studies in yeast further suggest that reduced expression of wild-type DNA polymerase epsilon may create a mutator phenotype with an increased probability of replication error (15). In addition, DNA polymerase delta, which replicates the lagging strand, has been shown to play a role in repairing polymerase epsilon errors (16).

In this study, we aimed to identify the mutational processes that may cause Isocitrate dehydrogenase 1 (*IDH1*) mutations, which are among the most recurrent mutations in cancer and occur in multiple tumor types including glioma, cholangiocarcinoma, and acute myeloid leukemia (17). IDH1 is a citric acid cycle enzyme that normally generates α-keto-glutarate. In cancer, gain-of-function *IDH1* mutations lead to production of the oncometabolite 2-hydroxyglutarate (2-HG) (17, 18). Since 2-HG cannot be used in major metabolic pathways, it accumulates in *IDH1*-mutated tumor cells and causes metabolic, epigenetic, and signaling alterations that promote tumorigenesis and allow immune evasion (17, 19–21). For example, 2-HG-induced global hypermethylation may block differentiation of *IDH1*-mutated tumor cells (19, 21). *IDH1* mutations occur early in tumorigenesis and are therapeutically targeted driver mutations (17, 22). Small-molecule inhibitors targeting mutated IDH1 protein, currently used in clinical practice, include Ivosidenib, Olutasidenib, and Vorasidenib (targets mutated IDH1/2) (22–29). Given the biological and clinical significance of *IDH1* mutations, it is important to understand their molecular etiology. Here, we evaluated whether recurrent *IDH1* mutations are aligned with known mutational signatures. Based on *in vitro* and computational analyses, we propose that *IDH1* R132C and R132G likely arise from APOBEC3-mediated deamination, while *IDH1* R132H may result from polymerase epsilon error.

## Materials and Methods

### Analysis of TCGA Data

Mutation, gene expression, methylation, and clinical data for tumors in the TCGA PanCancer dataset were accessed through cBioPortal (30–32). Gene expression data for adjacent normal bile duct tissue were accessed through UCSC Xena; gene expression data for adjacent normal tissue samples were not available for the other tumor types included in this study (33). Data were accessed in 2025, and data visualization and statistical analysis were completed using GraphPad Prism.

### Analysis of Replication Fork Directionality using OK-seq Data

Replication fork directionality (RFD) at the *IDH1* locus was analyzed using OK-seq data from HeLa S3 cells obtained from GEO (GSE193547). Genome-wide continuous RFD signals were processed with the OKseqHMM pipeline and aligned to the human reference genome hg19, using the bigWig file GSM5814025_Hela_S3_all_cutadapt_sort_delDupl_merge_hmm_RFD_cutoff10_bs1kb_sort.bw. RFD values across the *IDH1* locus (chr2:209,098,953–209,121,795) were extracted from the bigWig file using a custom Python workflow. Data extraction was performed with pyBigWig (v0.3.23), and downstream processing was carried out in pandas (v2.0.3). Visualization was generated in Matplotlib (v3.7.2) using the midpoint of each genomic interval to plot a continuous RFD profile. For genome-browser visualization, the RFD bigWig track was loaded into Integrative Genomics Viewer (IGV; v2.17.4).

### Cell culture

Human embryonic kidney 293FT cell line (R70007) was purchased from Thermo Fisher Scientific (Waltham, MA, USA). The human bladder cancer cell line HT-1376 (CRL-1472) was obtained from the American Type Culture Collection (ATCC; Manassas, VA, USA). Both cell lines were authenticated by the supplier and used within 2 years of purchase. These cell lines are listed in Cellosaurus as HEK293-FT (RRID: CVCL_6911) and HT-1376 (RRID: CVCL_1292). All cell lines were routinely tested for mycoplasma contamination using the MycoAlert® Mycoplasma Detection Kit (Lonza; LT07-318). Cells were maintained at 37 °C in a humidified incubator with 5% CO₂. HT-1376 cells were cultured in Eagle’s Minimum Essential Medium (EMEM; ATCC 30-2003) supplemented with 10% fetal bovine serum (FBS). 293FT cells were cultured in high-glucose Dulbecco’s Modified Eagle Medium (DMEM) supplemented with 10% FBS.

### *In Vitro* APOBEC3A DNA Cytosine Deamination Assays

Deamination assays were performed using APOBEC3A as previously described, with minor modifications (34). First, an open reading frame plasmid encoding Myc-tagged human *APOBEC3A* (Origene #RC220995) was transfected into 293FT cells (ThermoFisher Scientific #R70007) or HT1376 cells (ATCC #CRL-1472) grown in 10 cm plates. Following two days of incubation, APOBEC3A protein was isolated using the c-Myc tagged protein MILD PURIFICATION KIT Ver.2 (MBL Life Science #3305). For the *in vitro* deamination assays, partially purified APOBEC3A protein (0.25 µg) was incubated for one hour (APOBEC3A from 293FT cells) or 30 minutes (APOBEC3A from HT1376 cells) at 37°C in 10 µl reaction mixtures consisting of the following components: 20 pmol of custom DNA oligonucleotide probe (synthesized by IDT), 10 mM Tris-HCl (pH 8.0), 50 mM NaCl, and 1 mM DTT. Next, 5 µl of uracil DNA glycosylase (UDG) master mix, containing 0.75 units of UDG, was added to each reaction and incubated at 37°C for 40 minutes. Subsequently, 5 µl of 0.6 N NaOH was added to each tube and incubated at 37°C for an additional 20 minutes. An equal volume (20 µl) of 2x TBE-Urea Sample Buffer (Invitrogen) was then added, and the samples were heated at 95°C for 3 minutes, then cooled on ice for 5 minutes. DNA cleavage was analyzed using a 15% TBE-Urea polyacrylamide gel (Invitrogen), resolved at 100 V for 2 hours at room temperature in 1x TBE buffer. The gel images were captured using the Li-COR Odyssey imaging system with Alexa 488 fluorescence settings.

The custom oligonucleotide probes were single-stranded DNA sequences centered at the cytosines mutated in *IDH1* R132C and IDH1 R132H. Two sets of probes for each mutation were used: “a probes” contain cytosines within five nucleotides of the central cytosine of interest replaced with guanosines to avoid non-specific deamination, and “b probes” are wild-type sequences.

Positive control: 5’-6FAM-ATGATTATTATT**C**CCAATTATTTGT-3’

R132C-a (minus strand): 5’-6FAM-ATCATGATAGGT**C**GT***G***ATGCTTATG-3’

R132C-b (minus strand): 5’-6FAM-ATCATCATAGGT**C**GTCATGCTTATG-3’

R132H-a (plus strand): 5’-6FAM-CCATAAG***G***ATGA**C**GA***GG***TATGATGA-3’

R132H-b (plus strand): 5’-6FAM-CCATAAGCATGA**C**GACCTATGATGA-3’

### Analysis of CpG Methylation at the IDH1 R132H Locus

Methylated DNA immunoprecipitation (MeDIP) data for brain tissue were visualized in UCSC Genome Browser using the UCSF Brain DNA Methylation track (35, 36). Visualization focused on the CpG locus corresponding to the cytosine altered in *IDH1* R132H mutations on the plus strand of DNA. The genomic region was also evaluated for the presence or absence of CpG islands using the UCSC Genome browser CpG Island Track.

## Results

### Recurrent *IDH1* R132 Hotspot Mutations Involve Adjacent Cytosines on Opposite DNA Strands

*IDH1* mutations predominantly occur at amino acid position R132 and are recurrent in multiple cancer types including lower grade glioma (76.8%), cholangiocarcinoma (13.9%), acute myeloid leukemia (9.5%), glioblastoma (6.3%), and skin cutaneous melanoma (5.9%) (**Figure 1A**). *IDH1* R132H is most recurrent in lower grade glioma and glioblastoma, while *IDH1* R132C is the most common *IDH1* mutation in cholangiocarcinoma, acute myeloid leukemia, and melanoma. *IDH1* R132G, R132L, and R132S occur at lower frequencies. All five IDH1 R132 mutations arise from mutated cytosines at Chr2:209113112 or Chr2:209113113 (hg19), which are adjacent nucleotide positions on opposite DNA strands **(Figure 1B).**

**Figure 1:**
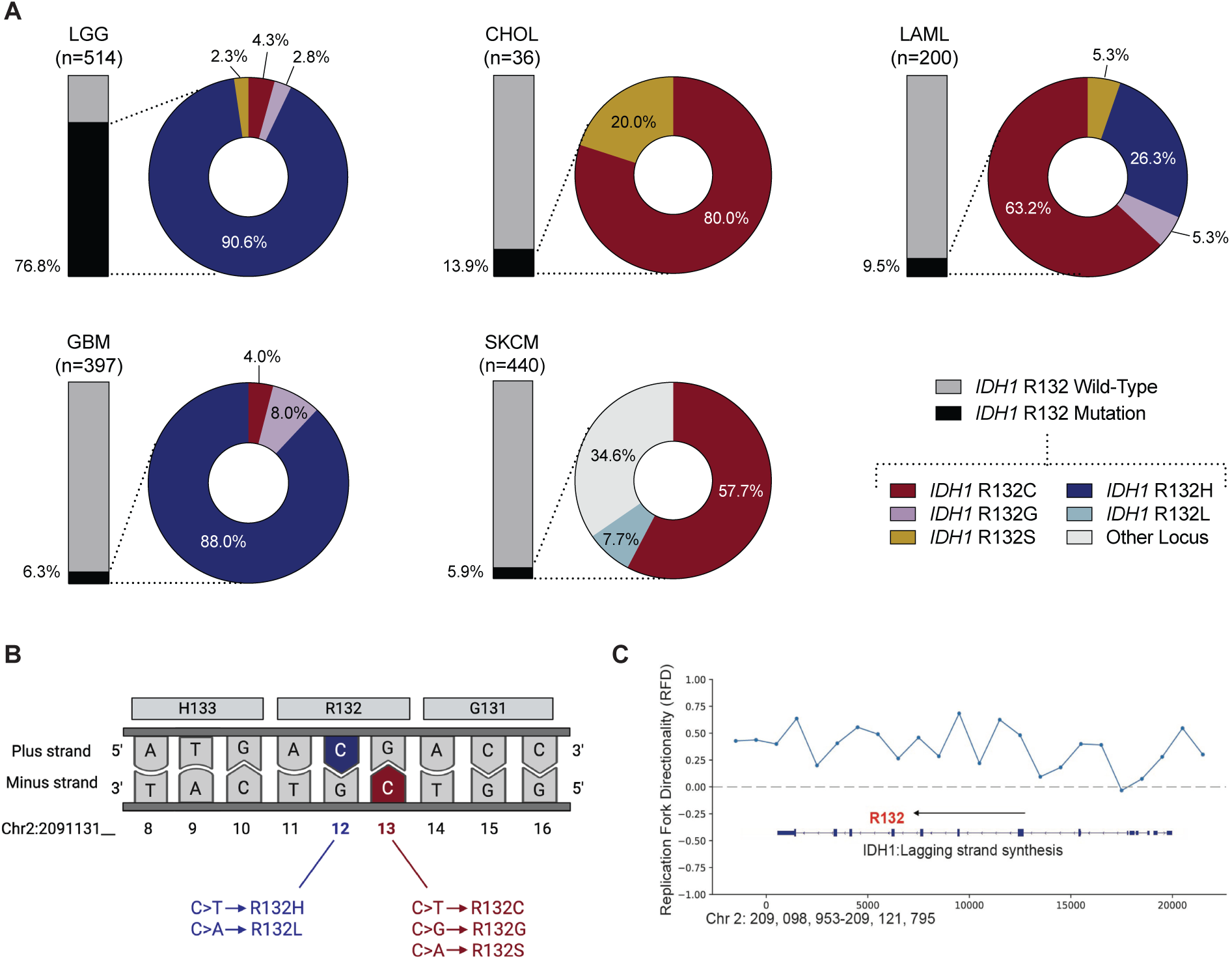
Recurrent *IDH1* R132 Mutations Map to Adjacent Cytosines on Opposite DNA Strands. **A:** *IDH1* mutation frequency in cancer types with at least 5% *IDH1* mutation rate in the TCGA PanCancer dataset: lower grade glioma (LGG), cholangiocarcinoma (CHOL), acute myeloid leukemia (LAML), glioblastoma multiforme (GBM), and skin cutaneous melanoma (SKCM). **B:** Sequence context and nucleotide substitutions for recurrent *IDH1* R132 mutations. **C:** DNA replication fork directionality (RFD) for *IDH1* in the context of the minus strand. Positive RFD values indicate lagging strand synthesis.

### *IDH1* R132C and R132G Likely Arise from APOBEC3-Mediated Mutagenesis

*IDH1* R132C and R132G are C>T and C>G mutations, respectively, occurring within a TpCpN motif, a sequence context compatible with APOBEC3-mediated deamination and APOBEC3-associated mutational signatures SBS2 and SBS13 **(Figure 1B)** (11, 37). Analysis of replication fork directionality using OK-Seq data showed that the cytosine altered in *IDH1* R132C occurs on the lagging strand template, whereas the cytosine altered in *IDH1* R132H occurs on the leading-strand template **(Figure 1C).** The *IDH1* R132C/R132G site also occurs within a region capable of forming a hairpin loop (**Figure 2A**). Together, these features make the TpCpN mutation context a permissive substrate for APOBEC3A and APOBEC3B activity.

**Figure 2:**
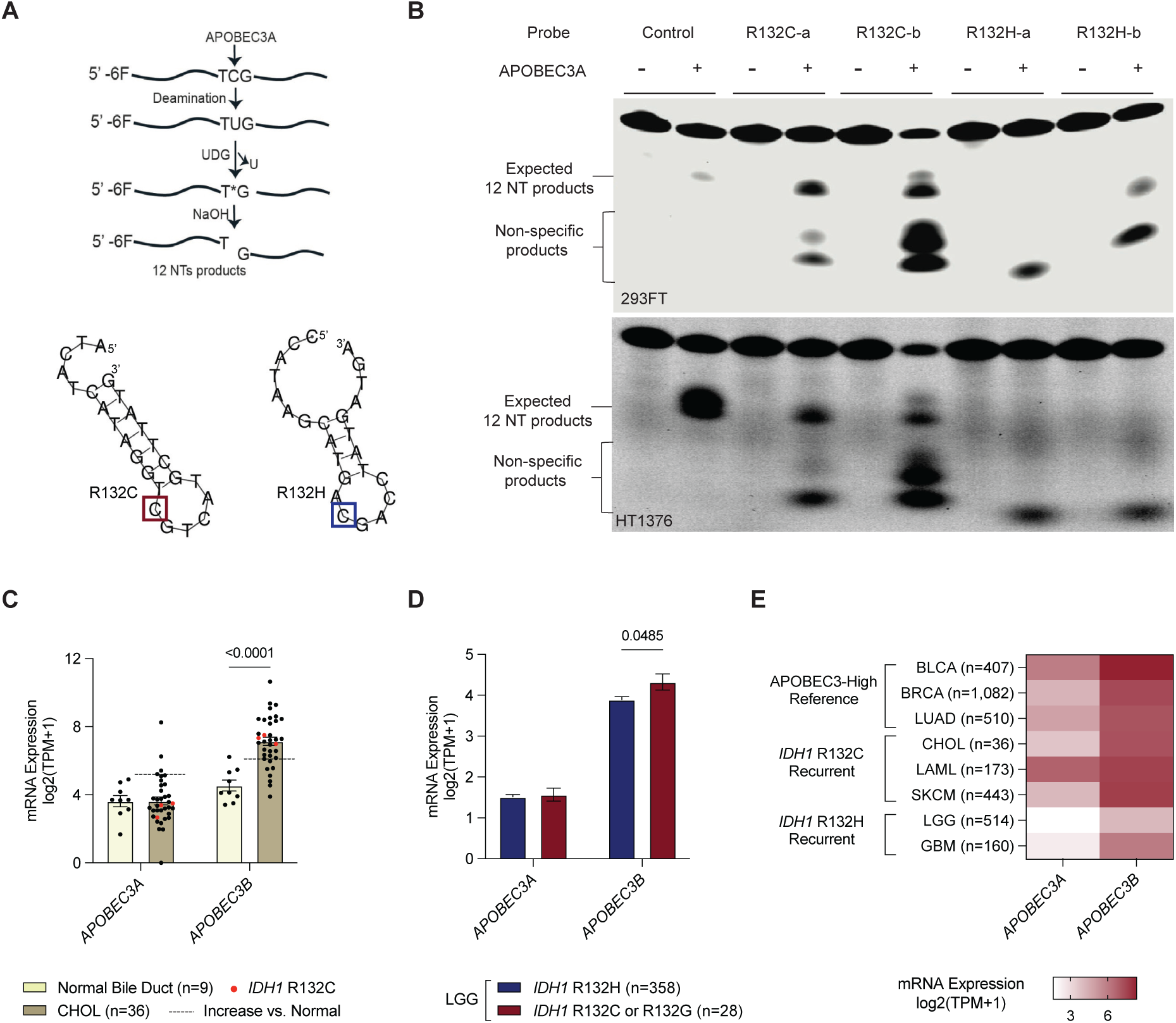
APOBEC3 Activity May Generate *IDH1* R132C and *IDH1* R132G. **A:** Schematic of *in vitro* deamination assay (upper), and visualization of DNA secondary structure for deamination assay probes (lower). Visualized probes are “b-probes,” which correspond to wild-type sequences centered at the recurrently mutated cytosines for *IDH1* R132C and *IDH1* R132H. Half-sized (12-nt) products indicate APOBEC3A-mediated deamination at the central cytosine of interest. **B:** *In vitro* deamination assays for recurrently mutated *IDH1* loci using APOBEC3A from 293FT cells (upper) or HT1376 cells (lower). **C:** *APOBEC3A* and *APOBEC3B* mRNA expression in normal bile duct tissue versus CHOL tumors. Tumors harboring *IDH1* R132C are indicated in red. Data are shown as mean and SEM. Thresholds for significant increases in *APOBEC3A* and *APOBEC3B* expression were calculated as the one-sided 95% confidence interval relative to normal bile duct tissue. P-values are for Welch’s t-tests. **D:** *APOBEC3A* and *APOBEC3B* expression in LGG tumors with *IDH1* R132C or R132G versus *IDH1* R132H. Data are shown as mean and SEM. P-values are for Welch’s t-tests. **E:** Heatmap showing mean APOBEC3A and APOBEC3B expression by cancer type, including those with predominant *IDH1* mutations and those known to have markedly high APOBEC3-associated mutational burdens.

To test this functionally, we performed in vitro deamination assays using APOBEC3A. APOBEC3A was selected as a representative APOBEC3 enzyme because it is more amenable than APOBEC3B to tagged affinity purification from human cell lysates (38), allowing in vitro assessment of substrate deamination with reduced confounding from endogenous APOBEC activity. APOBEC3A has also been shown to deaminate cytosines within a TCG context (37). Our assays demonstrated that cytosines within the *IDH1* R132C and R132G substitution contexts are deaminated in vitro (**Figure 2B**). In contrast, the cytosine altered in *IDH1* R132H was not subject to in vitro deamination, likely because it does not occur within the enzyme’s preferred TpC context (**Figures 2A and 2B**).

We next examined expression patterns of APOBEC3 family members in tumor tissues. We observed significant upregulation of *APOBEC3B* in cholangiocarcinoma—the tumor type in which *IDH1* R132C is most recurrent—compared with normal bile duct tissue (**Figure 2C**). In cholangiocarcinoma, *IDH1* R132C occurred exclusively in tumors with elevated *APOBEC3B* expression relative to normal tissue (**Figure 2C**). In lower-grade glioma, tumors harboring *IDH1* R132C exhibited higher A*POBEC3B* expression than tumors with *IDH1* R132H (**Figure 2D**).

Although *APOBEC3A* expression was not strongly elevated, it was clearly detectable in both normal and tumor tissues in cholangiocarcinoma (mean log2 TPM ∼3.8; **Figure 2C**) and higher than levels observed in lower-grade glioma (**Figure 2D**).

In a cross-cancer comparison, *APOBEC3A* and *APOBEC3B* expression were higher in tumor types with more recurrent *IDH1* R132C mutations—cholangiocarcinoma, acute myeloid leukemia, and melanoma—than in tumor types with more recurrent *IDH1* R132H mutations, including lower-grade glioma and glioblastoma (**Figure 2E**). *APOBEC3A* and *APOBEC3B* expression in tumor types with predominant *IDH1* R132C were also comparable to that observed in cancers known to have markedly high SBS2 and SBS13 mutational burdens, including breast cancer (**Figure 2E**) (39).

Tumors with *IDH1* R132C or R132G did not consistently exhibit higher *APOBEC3A,* or *APOBEC3B* expression compared with wild-type tumors across cancer types (**Supplementary Figures 1A**). This may partly reflect *IDH1* mutation–associated epigenetic remodeling, as increased methylation at cg22954818, located within a regulatory region upstream of the APOBEC3 gene cluster near the *APOBEC3A* promoter, was observed in mutant tumors compared with wild-type tumors (**Supplementary Figure 1B**).

Together with the sequence-context, strand-orientation, deamination, and expression analyses, the lack of an age association for *IDH1* R132C and R132G further supports an APOBEC3-related, rather than age-associated, mutational process (**Supplementary Figure 1C**).

### *IDH1* R132H May Arise from DNA Polymerase Epsilon Error

*IDH1* R132H is a CpG > TpG mutation within leading strand DNA, which raised the possibility that it arises from DNA polymerase epsilon error. Concordant with this suspected etiology, analysis of methylated DNA immunoprecipitation (Me-DIP) data revealed that the relevant cytosine is methylated in brain tissue—an epigenetic modification consistent with its location in a coding exon rather than a regulatory CpG island **(Figure 3A).** Methylation data from brain tissue was analyzed because *IDH1* R132H is most common in lower grade glioma and glioblastoma.

**Figure 3:**
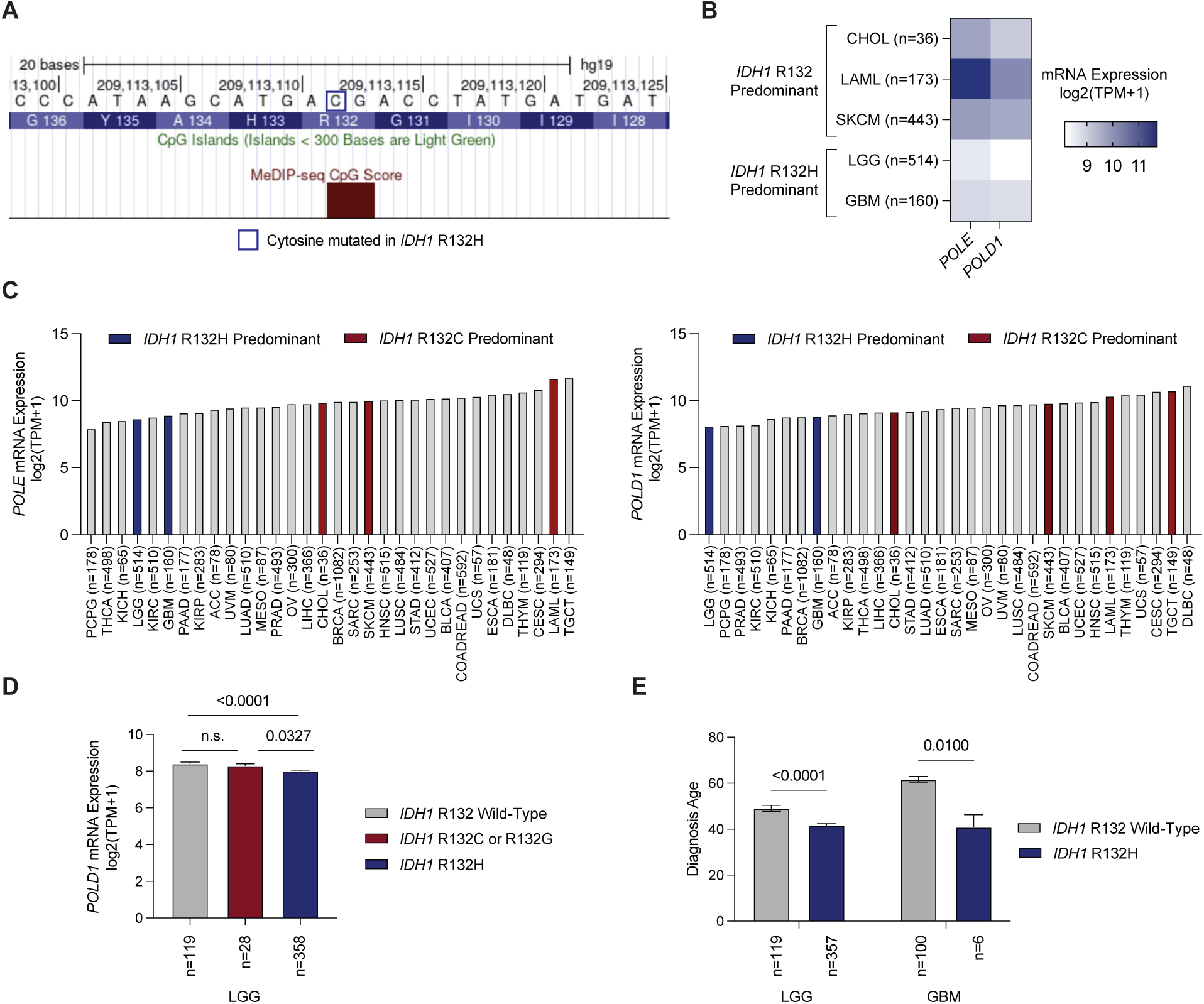
DNA Polymerase Epsilon Error is a Possible Etiology for *IDH1* R132H. **A:** Methylated DNA immunoprecipitation (Me-DIP) CpG score at the *IDH1* R132H mutation locus, visualized on the plus strand of DNA. A higher score indicates methylation. CpG islands, which were not present at the locus. **B:** Heatmap showing mean *POLE* and *POLD1* mRNA expression in cancer types with recurrent *IDH1* mutation. Cancer types are grouped by predominant *IDH1* mutation (R132C or R132H). **C:** *POLE* (left) and *POLD1* (right) mRNA expression in all tumor types. Cancer abbreviations follow standard TCGA nomenclature. Data are shown as means. **D:** *POLD1* mRNA expression in LGG tumors by *IDH1* mutation status (*IDH1* wild-type versus *IDH1* R132C or R132G versus *IDH1* R132H). Data are shown as mean and SEM. P-values are for post-hoc Dunn’s tests adjusted for multiple comparisons. **E:** Age at diagnosis for patients with *IDH1* wild-type versus *IDH1* R132H-mutated LGG or GBM tumors. Data are shown as mean and SEM. P-values are for Welch’s t-tests. Age data were not available for LAML.

Compared to tumor types with predominant *IDH1* R132C, lower grade glioma and glioblastoma had relatively low expression of *POLE* and *POLD1,* which encode the catalytic subunits for DNA polymerase epsilon and delta, respectively **(Figure 3B)**. Glioma and glioblastoma also had among the lowest *POLE* and *POLD1* expression levels across all tumor types, with lower grade glioma having the least *POLD1* expression of any cancer **(Figure 3C)**. Among lower grade gliomas, *IDH1* R132H-mutated tumors had significantly reduced *POLD1* expression compared to tumors with wild-type *IDH1*; *POLD1* expression was also reduced in *IDH1* R132H-mutated gliomas compared to *IDH1* R132C- or R132G-mutated gliomas **(Figure 3D)**. Taken together, these data suggest that DNA polymerase epsilon error in the setting of low POLE expression could lead to the initial genomic insult underlying *IDH1* R132H, while low *POLD1* expression may contribute to the mutation remaining unrepaired.

We also considered spontaneous deamination of 5-methylcytosine as an alternative cause of *IDH1* R132H, as this mutational process often causes CpG > TpG mutations of methylated cytosines. However, this mutational etiology is unlikely because it is age-associated unlike *IDH1* R132H mutations. In both lower grade glioma and glioblastoma, *IDH1* R132H was associated with younger age at diagnosis **(Figure 3E).**

## Discussion

Overall, our results implicate distinct mutational processes as likely sources of different oncogenic *IDH1* R132 alleles **(Figure 4)**. We first propose that APOBEC3 activity can generate *IDH1* R132C and R132G based on (i) concordance with APOBEC3A- and APOBEC3B ssDNA sequence motif contexts and lagging-strand features and (ii) direct *in vitro* evidence that APOBEC3A deaminates the relevant cytosine. The higher frequency of *IDH1* R132C relative to *IDH1* R132G may reflect the predominance of C>T over C>G outcomes following APOBEC3-mediated deamination and/or differences in oncogenic selection between the resulting amino acid substitutions (2, 7). C>A mutations can even more rarely arise from APOBEC3 activity, so it is possible that *IDH1* R132S also arises from APOBEC3-mediated deamination of the same cytosine (1, 7).

**Figure 4:**
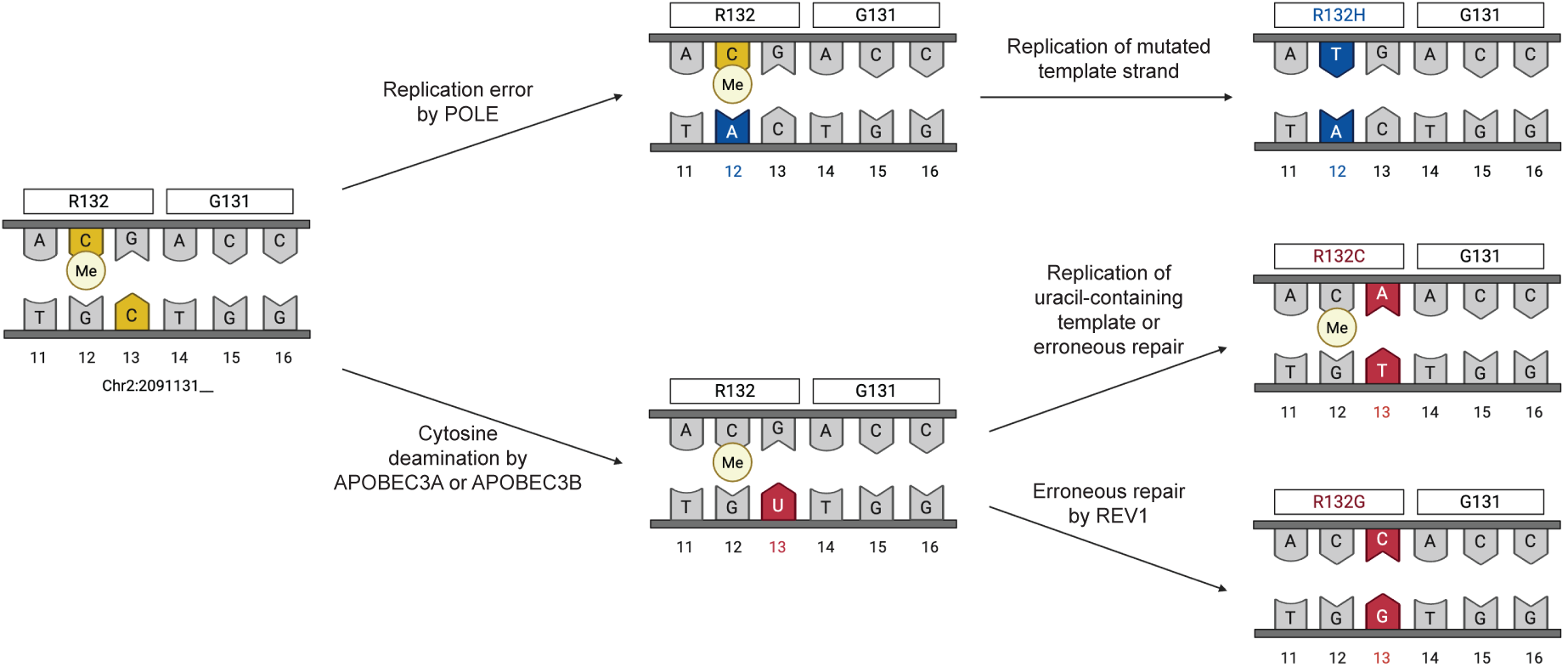
Summary of Likely Etiologies for *IDH1* R132C, R132G, and R132H Mutations. Different *IDH1* R132 mutations likely have distinct etiologies. DNA polymerase epsilon error during replication of leading-strand DNA may generate IDH1 R132H (CpG>TpG) mutations. In contrast, *IDH1* R132C (TpC>TpT) and R132G (TpC>TpG) may arise from APOBEC3A- and/or APOBEC3B-mediated deamination of lagging-strand DNA, generating an intermediate uracil-containing template; distinct replication or repair pathways can then generate C>T and C>G mutations.

Elevated *APOBEC3* expression in cholangiocarcinoma, acute myeloid leukemia, and melanoma plausibly explains why R132C is the most common *IDH1* mutation in these tumor types (1).

*IDH1* R132C and R132G are also common in chondrosarcoma, a cancer type that was not represented in the TCGA dataset (17). In cholangiocarcinoma, established risk factors—such as hepatitis infection and inflammatory conditions (e.g., primary sclerosing cholangitis)—could plausibly promote APOBEC3 induction, as *APOBEC3s* are interferon-stimulated genes upregulated by JAK-STAT and NF-κB signaling (2, 39). Interestingly, we did not observe *IDH1* R132C or R132G mutations in other tumor types with high APOBEC3 activity (e.g., bladder and breast cancers), suggesting tissue-specific differences in the selective advantage conferred by IDH1 R132 variants; similarly, other APOBEC3-associated hotspots, such as FGFR3 S249C, are highly tumor-type specific (10). Notably, the APOBEC3A- and APOBEC3B-associated signatures SBS2 and SBS13 have been documented in cholangiocarcinoma but are minimally present in glioma and glioblastoma, which much more frequently harbor R132H rather than R132C (1, 40). These patterns are consistent with our hypothesis that APOBEC3A and/or APOBEC3B may cause *IDH1* R132C and IDH1 R132G mutations in cholangiocarcinoma, while a different mutational process induces *IDH1* R132H in glioma and glioblastoma.

Finally, our data raise the possibility that *IDH1* R132H may result from DNA polymerase epsilon error, which is plausible given that the relevant cytosine occurs on the leading-strand template and is methylated. Comparatively low expression of the DNA polymerase epsilon catalytic-subunit gene, POLE, in lower-grade glioma and glioblastoma may help explain the predominance of *IDH1* R132H in these tumor types. It remains unclear whether low POLE expression emerges during tumorigenesis or reflects a baseline feature of healthy glial cells, as gene-expression data from adjacent normal tissue were not available for lower-grade glioma and glioblastoma. However, it is plausible that the minimal baseline turnover of healthy glial cells is accompanied by lower DNA polymerase epsilon expression, thereby creating a vulnerability to replication error during infrequent cell division events (e.g., reactive gliosis, NG2+ cell renewal) (41–44). This paradigm is consistent with *IDH1* R132 mutations arising early in tumorigenesis.

More broadly, multiple distinct mutational processes acting on the same codon to generate different amino acid substitutions likely reflects strong oncogenic selection pressure, underscoring the important role of *IDH1* R132 mutations in driving tumorigenesis across multiple cancer types.

### Conclusions

The molecular etiology of *IDH1* R132 mutations has remained unknown. Based on analysis of genomic features, gene expression data, and *in vitro* deamination assays, we propose that APOBEC3A and/or APOBEC3B activity may give rise to *IDH1* R132C and R132G mutations. In contrast, *IDH1* R132H may arise from replication error by DNA polymerase epsilon. These findings provide insight into the molecular oncogenesis of multiple cancer types, including cholangiocarcinoma and glioma.

## Data Availability Statement

Analyses used publicly available data from TCGA, Xena, and GEO.

## Author Contributions

K.E.B. and R.B. conceptualized the study design and wrote the manuscript. K.B. and E.U. completed bioinformatic analyses. B.L. conducted the *in vitro* deamination assays. All authors contributed to writing the materials and methods section and reviewed the final manuscript.

## Supplementary Figure

**Supplemental Figure 1:**
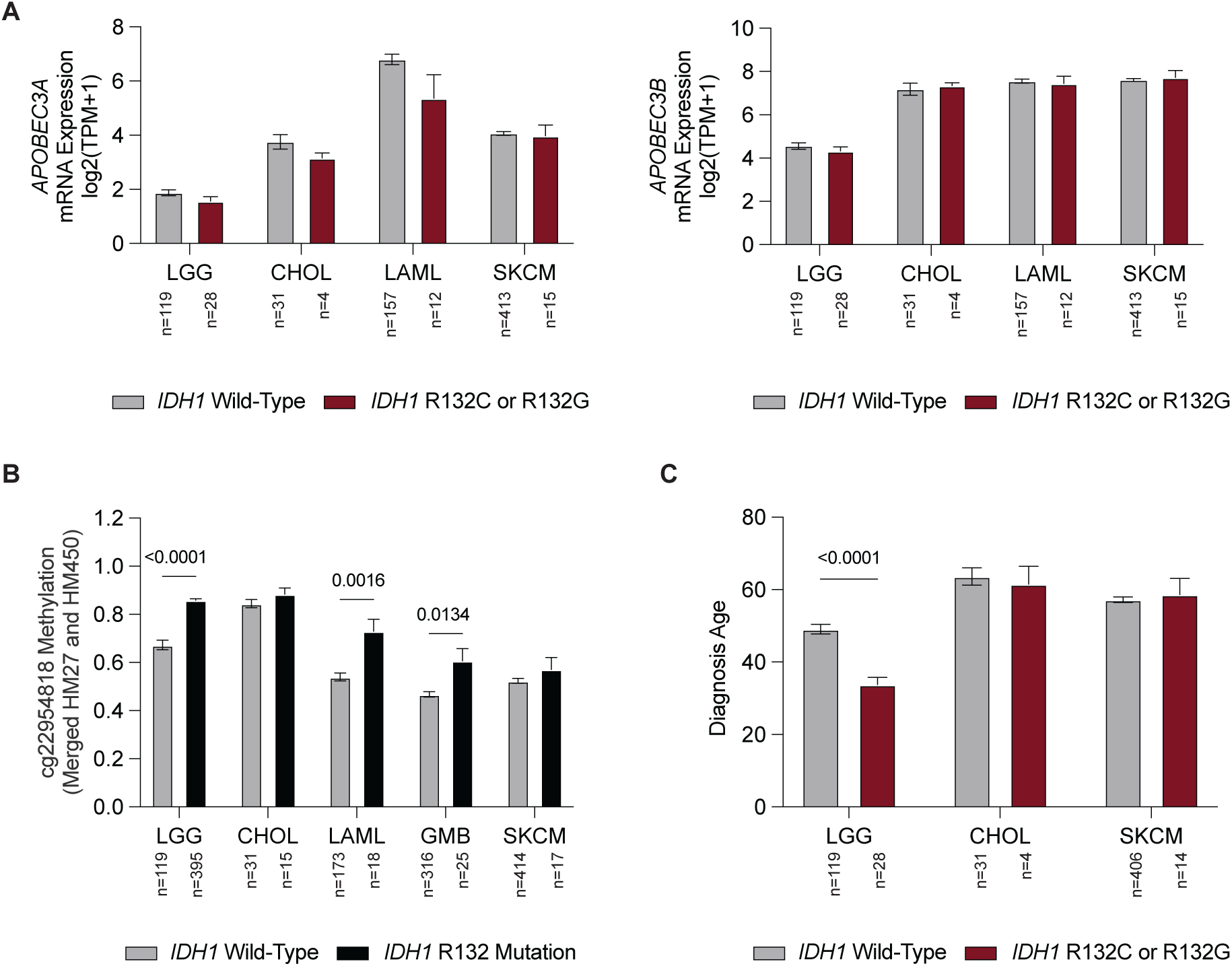
*APOBEC3* Expression, *APOBEC3* Methylation, and Patient Age at Diagnosis by *IDH1* Mutation Status. **A:** *APOBEC3A* and *APOBEC3B* expression in *IDH1* wild-type versus *IDH1* R132C- or R132G-mutated tumors by cancer type. Data are shown as mean and SEM. *P*-values are for Welch’s t tests. **B:** Methylation of cg22954818, which is within a regulatory region for *APOBEC3A,* in *IDH1* wild-type versus *IDH1* R132-mutated tumors. Tumors with any *IDH1* R132 mutation were included in the analysis because 2-HG-induced hypermethylation is not known to be allele-specific. Data are based on merged HM27 and HM450 Illumina microarrays. Data are shown as mean and SEM. *P*-values are for Welch’s t-tests. **C:** Age at diagnosis for patients with *IDH1* wild-type versus *IDH1* R132C- or *IDH1* R132G-mutated LGG, CHOL, or SKCM. Data are shown as mean and SEM. *P*-values are for Welch’s t tests. Age data were not completely available for LAML, or for GBM tumors with *IDH1* R132C or R132G mutations.

